# Alpha-actinin-1 stabilizes focal adhesions to facilitate sarcomere assembly in cardiac myocytes

**DOI:** 10.1101/2025.03.28.645933

**Authors:** James B Hayes, Anna M Bainbridge, Dylan T Burnette

## Abstract

Cardiac sarcomere assembly is a highly orchestrated process requiring integration between intracellular contractile components and extracellular adhesions. While α-actinin-2 (ACTN2) is well known for its structural role at Z-discs, the function of the “non-muscle” paralog α-actinin-1 (ACTN1) in cardiomyocytes remains unclear. Using human induced pluripotent stem cell-derived cardiac myocytes (hiCMs), we demonstrate that ACTN1 is essential for sarcomere assembly. siRNA-mediated depletion of ACTN1 disrupted Z-line formation and impaired sarcomere organization, defects that were rescued by exogenous ACTN1 but not ACTN2, revealing non-redundant functions. Unlike ACTN2, ACTN1 localized predominantly to focal adhesions and was required for adhesion maturation, as evidenced by reduced adhesion size and number following ACTN1 depletion. Live-cell imaging of vinculin dynamics showed decreased stability of adhesion-associated vinculin in ACTN1-deficient cells, whereas paxillin dynamics were unaffected. These results suggest that ACTN1 stabilizes focal adhesions to promote effective force transmission during sarcomere assembly.

## INTRODUCTION

Cardiac myocytes (CMs) are the beating muscle cells of the heart^1,2^. The fundamental contractile unit within beating CMs is the cardiac sarcomere^3^. Each cardiac sarcomere comprises overlapping filaments of actin and myosin II anchored between two Z-discs, which contain the actin cross-linking protein, α-actinin-2 (ACTN2)^4,5^. The process by which actomyosin, ACTN2, and other sarcomere components organize into ordered filaments within striated muscle cells is known conventionally as sarcomere assembly^6-9^.

ACTN2 is often called a “muscle-specific” α-actinin due to its high abundance in muscle^10^. Previous studies have demonstrated a requirement for ACTN2 during CM sarcomere assembly^9,11^. However, CMs also express ACTN1, a widely-expressed “non-muscle” α-actinin with unknown roles during assembly^12-14^. While studies of ACTN1 in CMs are limited, ACTN1 and ACTN2 have been shown to interact within CMs^15^, and another study showed that CMs upregulate *ACTN1* in response to cyclic mechanical stretch^16^. However, there have been no formal investigations into ACTN1 function within CMs, or within muscle cells in general, and our understanding of ACTN1 function therefore arises from studies of non-muscle cell lines^17-20^.

In non-muscle cells, ACTN1 serves prominent roles at focal adhesions, which mechanically anchor the cell to the external environment, the extracellular matrix (ECM)^17,21,22^. Focal adhesions are multi-protein complexes with different proteins appearing at different times, and contributing specific functions to the adhesion at large^23^. For instance, the adhesion protein paxillin is an early adhesion protein that serves as a scaffold between adhesions and the ECM^24^, while the protein vinculin serves specifically to link the adhesion complex to actomyosin networks^23,25,26^. In CMs, links between adhesions and actomyosin fulfill a required role during sarcomere assembly by coupling the intracellular contractile forces of assembly to the ECM^27^. For example, adhesion stability and size have been positively correlated with the capacity of sarcomere precursors to mature into sarcomeres^28^.

We have previously shown that sarcomere assembly can be studied *in vitro* using human cardiac myocytes derived from induced pluripotent stem cells (<u>**h**</u>uman <u>**i**</u>PSC-derived <u>**CM**</u>s, or hiCMs)^29^. Here, we present evidence within this model system that ACTN1 fulfills a required function during CM sarcomere assembly that is centered around adhesions. Following siRNA-mediated depletion of ACTN1, we find that hiCMs fail to assemble sarcomeres, and concomitantly that adhesions fail to enlarge during the normal period of sarcomere assembly. ACTN1 depletion impacted fluorescence lifetimes of vinculin, but not paxillin in adhesions, suggesting either a direct role for ACTN1 in linking pre-sarcomeric actomyosin to the adhesion complex, or an indirect role in stabilizing those links. Our data suggest that the role of ACTN1 at adhesions distinguishes it functionally from ACTN2 during sarcomere assembly in cardiac myocytes.

## RESULTS

ACTN1 and ACTN2 proteins exhibit high sequence and structural similarity^30^. Therefore, we first considered the hypothesis that ACTN1 and ACTN2 would have similar functions in CMs. ACTN2 is considered to be an essential component of the CM sarcomere and its depletion or knockout disrupts CM sarcomere assembly^9,11^. Our lab has demonstrated previously that CM sarcomere assembly can be expeditiously studied *in vitro* using <u>**h**</u>uman <u>**i**</u>PSC-derived <u>**c**</u>ardiac <u>**m**</u>yocytes (hiCMs) which, following a brief trypsinization period, assemble sarcomeres *de novo* within 24 hours of re-seeding onto glass coverslips^6^. This approach, when paired with siRNA-mediated protein depletion, can be used to test the essentiality of a protein or protein(s) for CM sarcomere assembly.

To determine if ACTN1 was essential for CM sarcomere assembly, we first asked if ACTN1 depletion from hiCMs prior to re-seeding would disrupt assembly. However, the assessment of ACTN1 depletion efficiency required anti-ACTN1 antibodies that were validated as being non-cross-reactive against the other ACTN paralogs expressed by hiCMs (ACTN2 and ACTN4). Therefore, we identified a commercially available anti-ACTN1 antibody (Invitrogen PA5-44889) raised against a divergent N-terminal sequence of ACTN1^30^. We then devised an experimental approach to validate the specificity of anti-ACTN1 by western blot, which separates proteins by molecular weight. Because the endogenous forms of ACTN1, ACTN2, and ACTN4 are indistinguishable by molecular weight, we introduced into hiCMs exogenous ACTN1-mEmerald, which we postulated would separate from endogenous bands due to the added molecular weight of mEmerald. We also introduced, into separate hiCM populations, either ACTN2-mEmerald or ACTN4-mEGFP, which would indicate cross-reactivity of anti-ACTN1 against ACTN2 and/or ACTN4. Finally, we included additional control populations where we either co-introduced ACTN1-mEmerald+ACTN4-mGFP (as a positive control) or introduced mEmerald only (as a negative control). We then collected whole cell lysates from each group for electrophoresis, membrane transfer, and western blotting with anti-ACTN1. The complete, uncropped western blot from this experiment is shown in **Fig. 1A**. While a visible band at the approximate size of endogenous alpha-actinin can be seen in all lanes, a larger band corresponding to exogenously-introduced protein is only visible in lanes containing lysate from hiCMs that expressed ACTN1-mEmerald **(Fig. 1A, lanes 1 and 4)**. From these data, we concluded that our anti-ACTN1 antibodies recognize ACTN1 and do not cross-react with ACTN2 or ACTN4.

**Figure 1:**
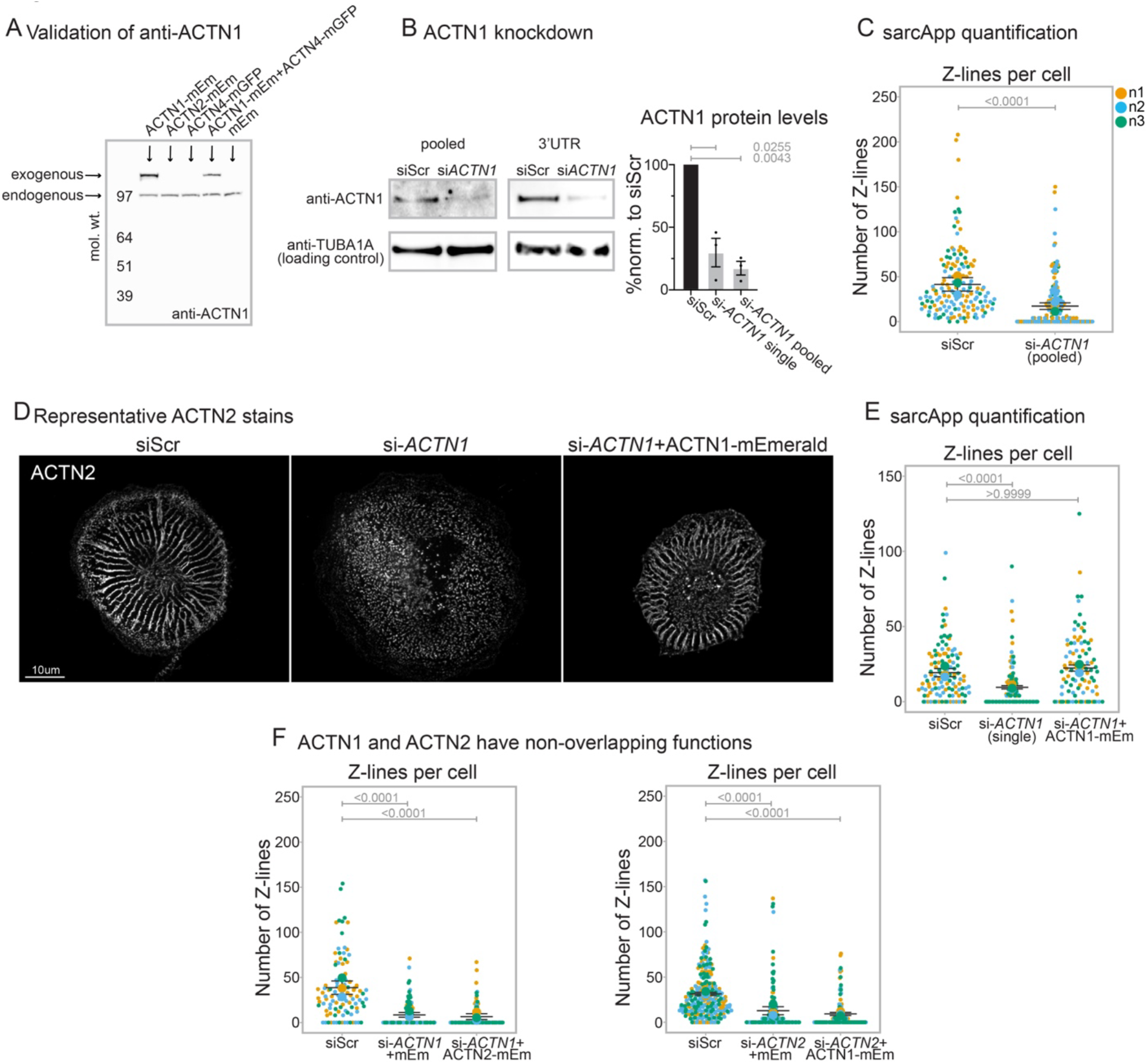
ACTN1 and ACTN2 exhibit non-redundant functions during sarcomere assembly in human iPSC-derived cardiac myocytes. **1A:** Western blot where each lane is loaded with whole cell lysate from hiCMs that were transfected with DNA encoding the indicated fluorescent fusion protein(s) for 24 hours. Blotted with anti-ACTN1 antibodies. **1B:** Left: crops from representative western blots using whole cell lysate from hiCMs following treatment with either non-targeting (si-Scr) or two different types of *ACTN1-*targeting (si-*ACTN1*) siRNAs. Top: blotted using anti-ACTN1 antibodies. Bottom: Same blot as top, stripped and re-blotted using anti-TUBA1A antibodies (tubulin; used as loading control). Right: ACTN1 protein levels in hiCMs (as a % of control) following depletion of ACTN1 using two different *ACTN1*-targeting siRNAs. **1C:** Z-lines per cell quantified in hiCMs treated with either non-targeting (si-Scr) or pooled *ACTN1-*targeting (si-*ACTN1* pooled) siRNAs. **1D:** ACTN2 stains in representative hiCMs following treatment with either non-targeting siRNA (si-Scr), 3’UTR-*ACTN1*-targeting siRNA (si-*ACTN1* single), or 3’UTR-*ACTN1-*targeting siRNA + exogenously expressed ACTN1-mEmerald (si-*ACTN1* + ACTN1-mEmerald). **1E:** Z-lines per cell as quantified by sarcApp for all imaged hiCMs across the treatment conditions from 1D. **1F:** Left: Z-lines per cell in hiCMs treated with either non-targeting siRNA (si-Scr), 3’UTR-*ACTN1*-targeting siRNA (si-*ACTN1*), or 3’UTR-*ACTN1-*targeting siRNA + exogenously expressed ACTN2-mEmerald (si-*ACTN1* + ACTN2-mEmerald). Right: Z-lines per cell in hiCMs treated with either non-targeting siRNA (si-Scr), 3’UTR-*ACTN2*-targeting siRNA (si-*ACTN2*), or 3’UTR-*ACTN2-*targeting siRNA + exogenously expressed ACTN1-mEmerald (si-*ACTN2* + ACTN1-mEmerald). **In panels C, E, and F:** small points represent individually-analyzed cells; large points represent population means of individually-performed biological replicates; black lines represent the mean (central line) and standard error (upper and lower) of the population means; color-code assigns individual data points and population means to associated biological replicate, represented by N1, 2, and 3 in panel C. **Statistical information:** B: Welch’s t-test. C: n=153 si-Scr, 165 si-*ACTN1* (pooled) cells, Mann-Whitney test; E: n=119 si-Scr, 98 si-*ACTN1* (single) cells, 94 si-*ACTN1* + ACTN1-mEmerald cells, Kruskal-Wallis multiple comparisons test; F: n=101 si-Scr cells, 83 si-*ACTN1 +* mEmerald cells, 90 si-*ACTN1+*ACTN2-mEmerald cells, Kruskal-Wallis multiple comparisons test; G: n=226 si-Scr cells, 146 si-*ACTN2 +* mEmerald cells, 119 si-*ACTN2+*ACTN1-mEmerald cells, Kruskal-Wallis multiple comparisons test.

Next, we assessed the efficiency of siRNA-mediated depletion of ACTN1 from hiCMs via western blot. We tested ACTN1 depletion capacity using two different approaches – a “pooled” approach (containing a multi-plex of four individual *ACTN1-*targeting siRNAs) and a “single” approach (containing a single, 3’-*ACTN1-*UTR-targeting siRNA). For both methods, our control groups were hiCMs exposed to media containing siRNA that was non-targeting (i.e., siScramble). After siRNA exposure, we collected whole cell lysates for subsequent electrophoresis, membrane transfer, and western blotting with anti-ACTN1 (to quantify ACTN1 levels), followed by anti-TUBA1A (as a loading control). Crops of representative western blots are shown in **Fig. 1B, left**. Across three experiments each, we observed that either approach (i.e., “pooled” or “single” *ACTN1-*targeting siRNA) resulted in significant reduction of ACTN1 protein levels from hiCMs compared to controls **(Fig. 1B, right)**. From these data, we concluded that *ACTN1-*targeting siRNAs can be used to deplete ACTN1 from hiCMs.

To test the hypothesis that ACTN1, like ACTN2, was essential for sarcomere assembly, we asked if depletion of ACTN1 from hiCMs prior to re-seeding would disrupt assembly. We treated individual hiCM populations with either siScramble or pooled *ACTN1-*targeting siRNAs, then trypsinized and re-seeded each group onto glass coverslips. We then chemically fixed hiCMs after a 24-hour period, a time-point in our assay by which the vast majority of re-seeded hiCMs have re-assembled sarcomeres^6^. To assess the impact of ACTN1 depletion on assembly, we labeled Z-lines (i.e., the 2-dimensional projection of Z-discs, which mark the sarcomere border) by immunofluorescence stains of ACTN2. We then quantified Z-lines per cell using sarcApp, our previously validated, automated analysis tool^31^. SarcApp analysis of three separate experiments (i.e., biological replicates) revealed that ACTN1-depleted hiCMs averaged fewer Z-lines per cell than control hiCMs **(Fig. 1C)**, indicating they had fewer sarcomeres. To determine if the defect in sarcomere assembly could be attributed specifically to reduced ACTN1 levels (and not siRNA-related off-target effects), we treated individual hiCM populations with either non-targeting siRNAs, 3’-*ACTN1-*UTR targeting siRNAs, or 3’-*ACTN1-*UTR targeting siRNAs with re-introduction of ACTN1 by transfection of plasmid encoding ACTN1-mEmerald, which lacks the siRNA-targeted 3’-*ACTN1-*UTR sequence. We then trypsinized, re-seeded, and fixed each hiCM population for subsequent ACTN2 staining and Z-lines quantification via sarcApp. Images of anti-ACTN2 stains within a representative hiCM from each population is shown in **Fig. 1D**. SarcApp analysis of three experiments revealed that ACTN1-depleted hiCMs averaged fewer Z-lines per cell than control hiCMs or ACTN1-depleted hiCMs expressing ACTN1-mEmerald **(Fig. 1E)**. Taken together, these data suggest ACTN1 has a required function during CM sarcomere assembly.

It could be inferred from the structural similarities of ACTN1 and ACTN2 that they would exhibit functional overlap during sarcomere assembly^30^. In such a scenario, the impact of ACTN1 depletion on assembly could be explained by the simple depletion of actin cross-linkers, and may be reversible through exogenous over-expression of ACTN2. Alternatively, if ACTN1 and ACTN2 have non-overlapping functions during assembly, then the over-expression of ACTN2 would be insufficient to restore Z-lines. Therefore, we treated hiCM populations with either non-targeting siRNAs, 3’-*ACTN1-*UTR targeting siRNAs + plasmid-encoded mEmerald, or 3’-*ACTN1-*UTR targeting siRNAs + plasmid-encoded ACTN2-mEmerald. We then trypsinized, re-seeded, and fixed each hiCM population for subsequent ACTN2 staining and Z-lines quantification via sarcApp. SarcApp analysis revealed that ACTN1-depleted hiCMs averaged fewer Z-lines than controls whether we introduced mEmerald or ACTN2-mEmerald **(Fig. 1F, left)**. These data suggested ACTN2 could not functionally compensate for the loss of ACTN1 during sarcomere assembly. Next, we performed the reciprocal experiment, asking if re-introduction of ACTN1 restored Z-lines in ACTN2-depleted hiCMs. We treated hiCM populations with either non-targeting siRNAs, *ACTN2-*targeting siRNAs + plasmid-encoded mEmerald, or *ACTN2-*targeting siRNAs + plasmid-encoded ACTN1-mEmerald. As in ACTN1-depleted hiCMs, ACTN2-depleted hiCMs had fewer Z-lines than controls whether we introduced mEmerald or ACTN1-mEmerald **(Fig. 1F, right)**. Taken together, these data suggest ACTN1 and ACTN2 have non-overlapping functions during CM sarcomere assembly.

If the functions of ACTN1 and ACTN2 in CMs are non-overlapping, we hypothesized they might exhibit unique localization patterns in hiCMs. When we simultaneously localized both proteins in hiCMs using immunofluorescence, anti-ACTN2 antibodies localized throughout hiCMs, with a classically prominent localization at Z-lines **(Fig. 2A, left)**. Conversely, anti-ACTN1 antibodies localized almost exclusively to structures resembling focal adhesions relatively little localization at Z-lines **(Fig. 2A, left)**. Co-immunofluorescence of ACTN1 and the adhesion protein paxillin revealed co-localization of ACTN1 with paxillin, confirming the enrichment of ACTN1 at adhesions **(Fig. 2A, right)**. These data are consistent with the known functions of ACTN1 at non-muscle cell adhesions^17,18,21,22^.

**Figure 2:**
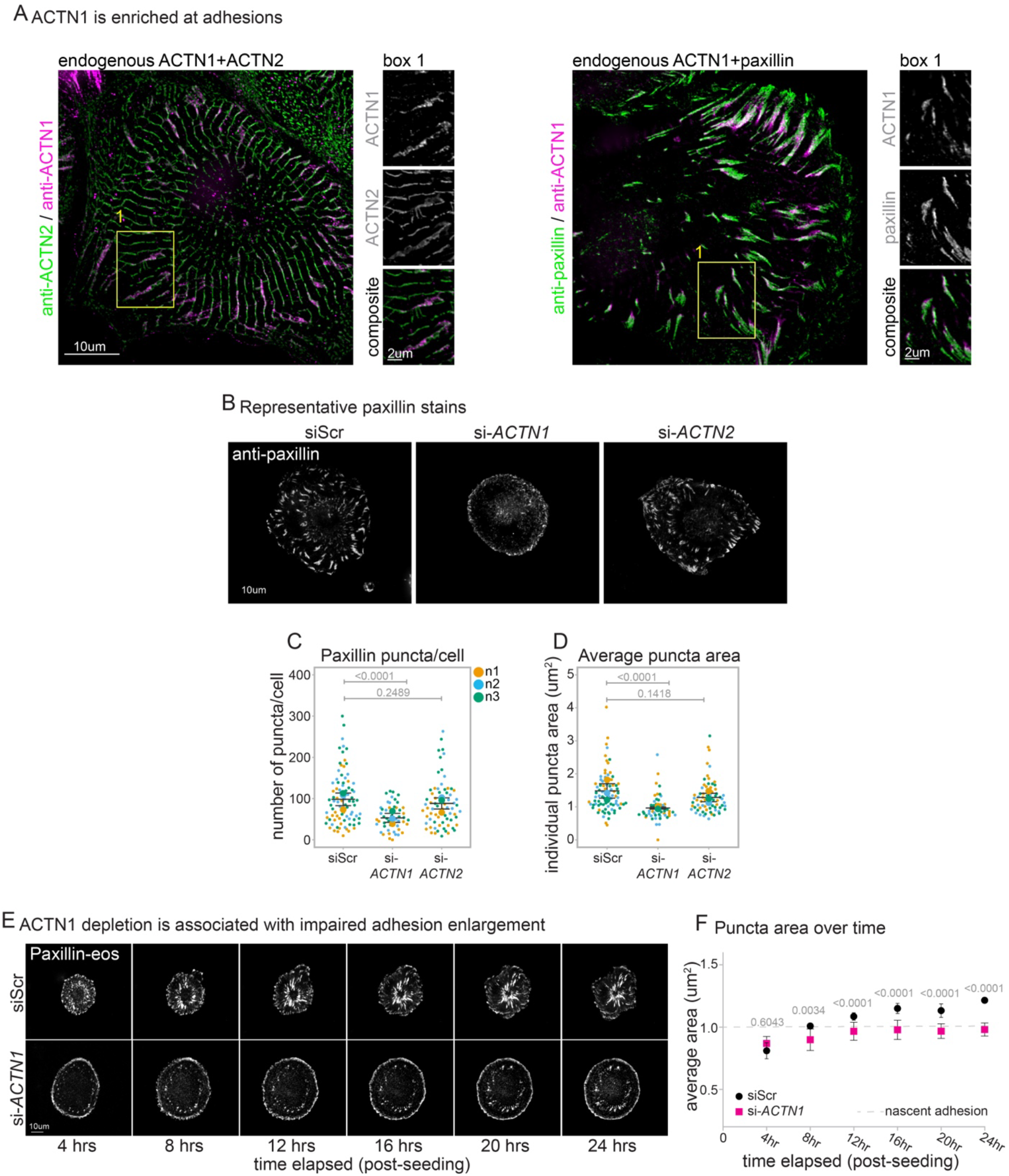
ACTN1 but not ACTN2 is required for adhesion maturation during hiCM sarcomere assembly. **2A:** Left: Immunofluorescence co-stains of endogenous ACTN2 and ACTN1 in a human iPSC-derived cardiac myocyte (hiCM) using anti-ACTN1 and anti-ACTN2 antibodies. Right: immunofluorescence co-stains of endogenous ACTN1 and paxillin in a hiCM using anti-ACTN1 and anti-paxillin antibodies. **2B:** Paxillin stains in representative hiCMs following treatment with either non-targeting siRNA (si-Scr), *ACTN1-*targeting siRNA (si-*ACTN1*), or *ACTN2-*targeting siRNA (si-*ACTN2*). **2C:** Number of individual paxillin puncta per cell in hiCMs following treatment with either non-targeting siRNA (si-Scr), *ACTN1-*targeting siRNA (si-*ACTN1*), or *ACTN2-*targeting siRNA (si-*ACTN2*). **2D:** Average size of all individual puncta per cell in hiCMs following treatment with either non-targeting siRNA (si-Scr), *ACTN1-*targeting siRNA (si-*ACTN1*), or *ACTN2-*targeting siRNA (si-*ACTN2*). **2E:** Montages of paxillin-eos green fluorescence in a representative hiCM following treatment with either non-targeting (si-Scr) or *ACTN1-*targeting (si-*ACTN1*) siRNA. **2F:** Average area of all paxillin-eos puncta in hiCMs across time following treatment with either non-targeting (si-Scr) or *ACTN1-*targeting (si-*ACTN1*) siRNA. Points (circles/boxes) represent mean of population means; error bars represent standard error of the mean. **In panels C and D:** small points represent individually-analyzed cells; large points represent population means of individually-performed biological replicates; black lines represent the mean (central line) and standard error (upper and lower) of the population means; color-code assigns individual data points and population means to associated biological replicate, represented by N1, 2, and 3 in panel C. **Statistical information:** C: 84 siScr cells, 50 si-*ACTN1* cells, 71 si-*ACTN2* cells, ordinary one-way ANOVA; D: 84 siScr cells, 50 si-*ACTN1* cells, 71 si-*ACTN2* cells, ordinary one-way ANOVA; F: n=1,702 si-Scr adhesions and 666 si-*ACTN1* adhesions (4 hours), n=1,933 si-Scr adhesions and 724 si-*ACTN1* adhesions (8 hours), n=1,962 si-Scr adhesions and 854 si-*ACTN1* adhesions (12 hours), n=1,902 si-Scr adhesions and 923 si-*ACTN1* adhesions (16 hours), n=1,821 si-Scr adhesions and 872 si-*ACTN1* adhesions (20 hours), n=1,750 si-Scr adhesions and 1,102 si-*ACTN1* adhesions (24 hours), all tests are Welch’s t-test.

To test if either ACTN1 or ACTN2 is required for CM adhesion formation, we treated hiCMs with non-targeting, *ACTN1*-targeting (si-*ACTN1*), or *ACTN2-*targeting (si-*ACTN2*) siRNAs, then trypsinized, re-seeded, and fixed hiCMs after 24 hours. We then stained hiCMs using antibodies against paxillin, which localizes to adhesions, and quantified the number and area of individual anti-paxillin puncta. Images of anti-paxillin stains within a representative hiCM from each population is shown in **Fig. 2B**. Across three experiments, ACTN2-depleted and control hiCMs had similar numbers of puncta/cell and average puncta sizes **(Fig. 2C-D)**. Meanwhile, ACTN1-depleted hiCMs averaged fewer puncta/cell and puncta were smaller **(Fig. 2C-D)**. Taken together, these data suggest ACTN1 but not ACTN2 functions in CMs to regulate the size and/or the number of CM adhesions.

In CMs, adhesions and sarcomeres are directly physically coupled^28^. Prior to sarcomere assembly, nascent (i.e., immature) adhesions link the structures which serve as sarcomere precursors to the ECM^28^. As sarcomere assembly proceeds, increased contractile forces within the cell drive increased biophysical demands on cell-ECM contact sites. To meet those demands, adhesions must undergo a maturation process that characteristically results in enlargement of the adhesion^32,33^. Our group has shown previously that adhesion enlargement is required for hiCM sarcomere assembly to proceed and that adhesion size directly correlates with adhesion maturity^28^. Therefore, we asked if the defect in sarcomere assembly in ACTN1-depleted hiCMs could be attributed to a defect in adhesion maturation. If so, we would expect to observe that adhesions fail to enlarge during the specific time window that sarcomere assembly occurs. In our hands, re-seeded hiCMs assemble the majority of their sarcomeres between ∼6 hours and 24 hours post re-seeding, with most hiCMs being devoid of sarcomeres prior to the 6-hour mark^6^. Therefore, we treated hiCMs with either non-targeting or si-*ACTN1-*targeting siRNAs, then introduced a plasmid encoding paxillin-eos prior to re-seeding, enabling us to track the progression of individual adhesion plaques in live hiCMs over time. Montages of paxillin-eos are shown within a representative hiCM from each population in **Fig. 2E**. We measured the progression of adhesion area in the same cells from the period of 4 hours to 24 hours post-seeding. We found that, at 4 hours post-seeding, ACTN1-depleted and control (siScr) hiCMs exhibited no differences in average adhesion area **(Fig. 2F)**. By 8 hours post-seeding, average adhesion area in control hiCMs had begun to increase, a trend which generally continued until the 24-hour mark and culminated in an average area of 1.2um^2^ **(Fig. 2F)**. Meanwhile, average adhesion area in ACTN1-depleted hiCMs increased more slowly and never at any point eclipsed 1.0um^2^, the generally-accepted cutoff of nascent adhesions^34^. These data suggest that nascent adhesions fail to mature in CMs depleted of ACTN1.

Adhesion maturation requires tension development through the coupling of the adhesion complex with contractile actomyosin inside the cell^35-37^. Therefore, we hypothesized that ACTN1 stabilizes the link between adhesions and the actomyosin of CM sarcomere precursors during CM sarcomere assembly. The adhesion protein vinculin canonically links adhesion complexes directly to actomyosin^23,25,26^ and depletion of vinculin from hiCMs prior to re-seeding disrupts sarcomere assembly^28^. Therefore, we treated hiCMs with either non-targeting or si-*ACTN1-*targeting siRNAs, then introduced a plasmid encoding vinculin-eos prior to re-seeding. Vinculin-eos is a photo-convertible fluorescent protein, enabling us to convert a specific subpopulation to emit red light (instead of green) and track that red-emitting population over time. Furthermore, a measure of vinculin stability at the adhesion can be ascertained from the rate of red fluorescence loss. Thus, we converted a subpopulation of vinculin-eos from green to red and measured red fluorescence intensity over time in control and ACTN1-depleted hiCMs, beginning at 18 hours post-seeding. Montages showing fluorescence of photo-converted vinculin-eos over time are shown within a representative hiCM from each population in **Fig. 3A**. Shown in **Fig. 3B** are population-averaged (across three experiments) curves of vinculin-eos red fluorescence intensity over time, each normalized to maximum fluorescence intensity. From curve fits of individual cell curves, we quantified fluorescence loss rate on a per-cell basis using two separate methods, i) the fluorescence lifetime of photo-converted vinculin-eos at the adhesion (i.e., half-life) and ii) the fraction of photo-converted vinculin-eos signal which stayed bound at the adhesion (i.e, mobile fraction) during the period of imaging. Across three experiments, the average fluorescence half-life (in minutes) of photo-converted vinculin-eos was shorter in ACTN1-depleted hiCMs compared to controls **(Fig. 3C)**, suggesting ACTN1 depletion increased the rate at which vinculin-eos dissociated from the adhesion complex. Furthermore, the average mobile fractions of photo-converted vinculin-eos were increased in ACTN1-depleted hiCMs **(Fig. 3D)**, suggesting ACTN1 depletion decreased the fraction of vinculin-eos which stayed bound at the adhesion during the imaging period. Taken together, these data suggest ACTN1 is required for the stable association or the retention of vinculin at CM focal adhesions during periods of CM sarcomere assembly.

**Figure 3:**
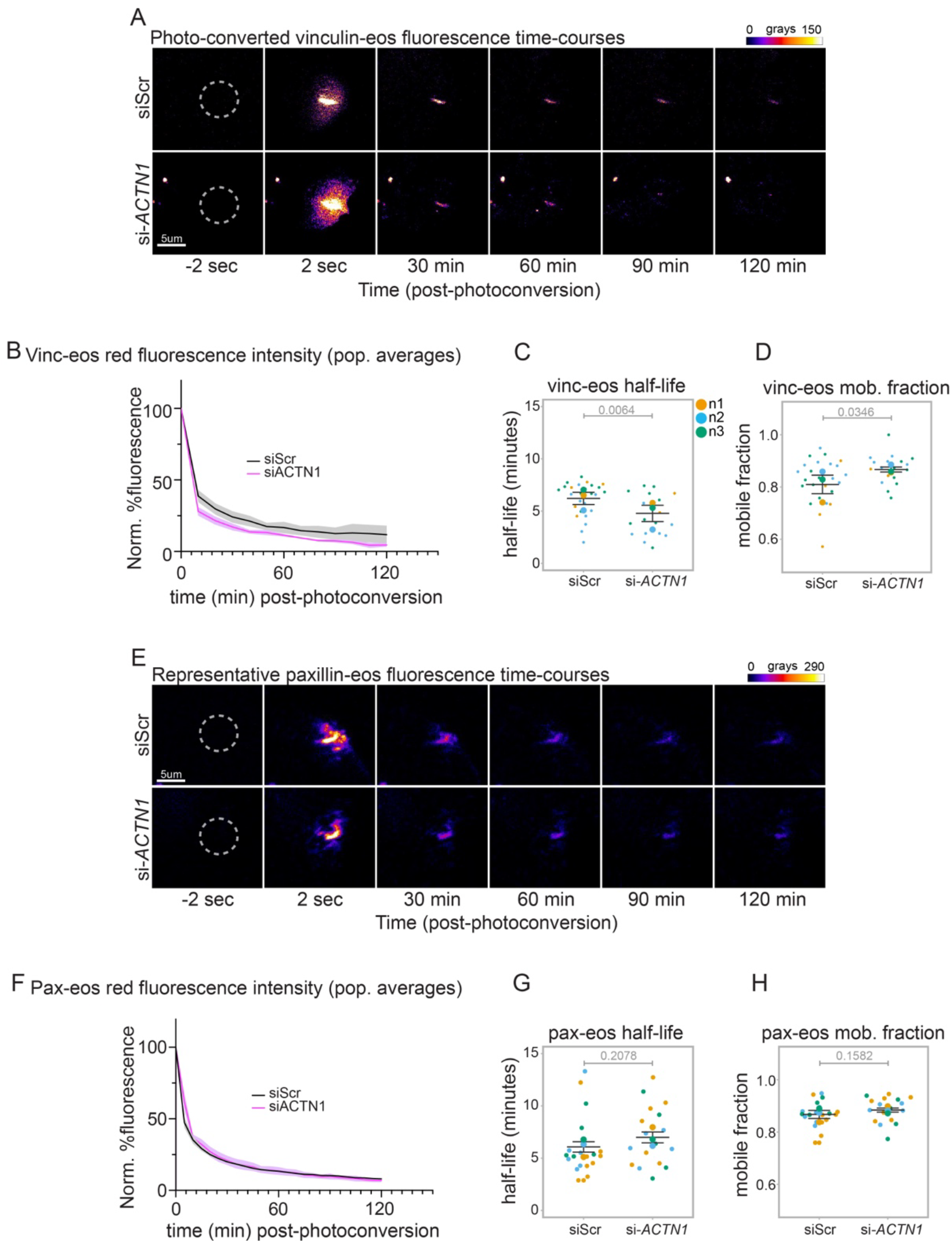
ACTN1 depletion impacts the turnover of vinculin but not paxillin within hiCM adhesions. **3A:** Montages of photo-converted vinculin-eos fluorescence in representative hiCMs following treatment with either non-targeting (si-Scr) or *ACTN1-*targeting (si-*ACTN1*) siRNAs. **3B:** Population-averaged curves of photo-converted vinculin-eos red fluorescence signal (normalized to initial/maximum photoconverted signal). Solid lines represent mean of population means; shaded regions represent standard error of the means. **3C:** Half-life of photo-converted vinculin-eos in hiCM adhesions following treatment with either non-targeting (si-Scr) or *ACTN1-*targeting (si-*ACTN1*) siRNAs, calculated from individual curves of photo-converted signal. **3D:** Mobile fraction of photo-converted vinculin-eos in hiCM adhesions following treatment with either non-targeting (si-Scr) or *ACTN1-*targeting (si-*ACTN1*) siRNAs, calculated from all individual curves of photo-converted signal. **3E:** Montages of photo-converted paxillin-eos fluorescence in representative hiCMs following treatment with either non-targeting (si-Scr) or *ACTN1-*targeting (si-*ACTN1*) siRNAs. **3F:** Population-averaged curves of photo-converted paxillin-eos red fluorescence signal (normalized to initial/maximum photoconverted signal). Solid lines represent mean of population means; shaded regions represent standard error of the means. **3G:** Half-life of photo-converted paxillin-eos in hiCM adhesions following treatment with either non-targeting (si-Scr) or *ACTN1-*targeting (si-*ACTN1*) siRNAs, calculated from individual curves of photo-converted signal. **3H:** Mobile fraction of photo-converted paxillin-eos in hiCM adhesions following treatment with either non-targeting (si-Scr) or *ACTN1-*targeting (si-*ACTN1*) siRNAs, calculated from all individual curves of photo-converted signal. **In panels C, D, G, and H:** small points represent adhesions in individually-analyzed cells; large points represent population means of individually-performed biological replicates; black lines represent the mean (central line) and standard error (upper and lower) of the population means; color-code assigns individual data points and population means to associated biological replicate, represented by N1, 2, and 3 in panel C. **Statistical information:** C-D: n=26 siScr adhesions, 18 si-*ACTN1* adhesions, Welch’s t-test; D: G-H: n=21 siScr adhesions, 17 si-*ACTN1* adhesions, Welch’s t-test.

As our data suggested that ACTN1 regulated the turnover of the adhesion protein vinculin, we next hypothesized that ACTN1 may regulate turnover of the entire adhesion complex. We therefore asked if ACTN1 depletion would likewise impact turnover of other adhesion proteins. We had observed earlier that ACTN1-depleted hiCMs exhibited fewer anti-paxillin puncta, suggesting it may also influence paxillin stability. Therefore, we treated hiCMs with either non-targeting or si-*ACTN1-*targeting siRNAs, then introduced a plasmid encoding paxillin-eos prior to re-seeding. To measure paxillin stability at the adhesion, we converted a subpopulation of paxillin-eos from green to red and measured red fluorescence intensity over time in control and ACTN1-depleted hiCMs. Montages showing fluorescence of photo-converted paxillin-eos over time are shown within a representative hiCM from each population in **Fig. 3E**. Shown in **Fig. 3F** are population-averaged (across three experiments) curves of paxillin-eos red fluorescence intensity over time, each normalized to maximum fluorescence intensity. Similar to vinculin-eos, we quantified from single-cell curve-fits the i) half-life and the ii) mobile fraction of photo-converted paxillin-eos signal. Across three experiments, the half-lives (in minutes) and mobile fractions of photo-converted paxillin-eos were no different in ACTN1-depleted hiCMs and controls **(Fig. 3G-H)**. These data suggest ACTN1 does not impact the stability of paxillin at CM adhesions. Taken together with results from **Fig. 3A-D**, our results suggest ACTN1 regulates the stability of proteins required for CM adhesion maturation but not those involved in adhesion formation.

## DISCUSSION

We provide in the current study evidence of a required function during CM sarcomere assembly for ACTN1, a widely-expressed, “non-muscle” paralog of muscle-specific ACTN2. Although ACTN1 and ACTN2 each appeared to localize at CM adhesions, only ACTN1 depletion produced a measurable impact on adhesion morphology. We observe that ACTN1 depletion disrupts the progressive increase in adhesion size normally seen in hiCMs that are in a period of active sarcomere assembly. The failure of adhesions in ACTN1-depleted hiCMs to increase in size suggests a defect in force transmission through the adhesion, which can also explain the defect in sarcomere assembly. The defect in assembly was rescued by the re-introduction of exogenous ACTN1 but not by the exogenous introduction of ACTN2, supporting the conclusion that the defect is not attributable to a general reduction in actin cross-linking activity but instead to the removal of some required function that is specifically fulfilled by ACTN1.

Our data support a model whereby ACTN1 is required for adhesion maturation in hiCMs. We observed that ACTN1-depleted hiCMs contained smaller adhesions than controls; however, they also contained fewer adhesions, suggesting some role for ACTN1 in adhesion formation. We suggest that hiCMs may form adhesions during periods of sarcomere assembly as part of a two-tiered process – an initial wave of nascent adhesions forms in the absence of or under only low actomyosin contractile forces (similarly to in non-muscle cells^38^). The ACTN1-dependent maturation of these initial adhesions may promote subsequently the formation of additional nascent adhesions. Such a model accounts for each of the phenotypes we observe in ACTN1-depleted hiCMs.

Our conclusion that ACTN1 and ACTN2 have non-redundant functions in CMs is in accordance with numerous studies in non-muscle cells, which similarly reported non-redundant functions for distinct paralogs of α-actinin^39-41^. In CMs, recent reports have linked ACTN1 but not ACTN2 as being responsive to cycles of mechanical stretch^15^. This study and our own support largely the conclusion that ACTN1 and ACTN2 in CMs have functionally diverged into specialized roles. Our results may be consistent with a role for ACTN1 during a CM mechanical stretch response, as adhesions of CMs in 2-dimensional culture (as in the current study) are held as structural analogs to the striated-muscle costamere, which has well-known roles in stretch-sensing^42-44^. Thus, ACTN1 functioning at adhesions should not conceptually limit it to the process of sarcomere assembly.

It is interesting to consider how adhesions would mature in ACTN2-depleted hiCMs but not ACTN1-depleted hiCMs. Adhesions in ACTN2-depleted hiCMs were indistinguishable from controls. This observation could be interpreted to mean that ACTN1 functions sequentially upstream of ACTN2 during sarcomere assembly, though ACTN2 is still required for Z-line formation^9^. Consistent with such a role for ACTN1, previous models of CM sarcomere assembly have postulated that other “non-muscle” paralogs of muscle-specific proteins serve upstream roles with respect to their muscle paralogs. For instance, a “non-muscle” paralog of muscle myosin II, myosin IIB, is well-established as a required component for sarcomere assembly, and is thought to function upstream of muscle myosin IIs^6-8^. ACTN1 appears to fit within this paradigm as another “non-muscle” protein with a required role that precedes, in the sequence of assembly, that of its muscle paralog, ACTN2.

We observe that ACTN1 depletion decreased life-times and increased mobile fractions of photo-converted vinculin, but not paxillin at adhesions. These data suggest ACTN1 does not necessarily stabilize the adhesion at large, or play a general scaffolding role, but instead plays a more precise role centered around force transmission (in line with our other data) and/or specifically linking the adhesion to actomyosin. In non-muscle cells, vinculin canonically serves this role through a conformational shift that might require stable association with the adhesion complex^45^. However, it is also possible that ACTN1 itself must be present to form proper cross-links with pre-sarcomeric actomyosin, as different actinin paralogs are known to uniquely cross-link actin networks^46^. Disruption of the link between the adhesion and the pre-sarcomeric machinery through ACTN1 depletion may indirectly influence vinculin lifetimes through the inability of the cell to generate tension on the substrate^28^.

Here, we add another widely-expressed version of a muscle-specific sarcomere protein to the molecular parts list of proteins required for CM sarcomere assembly. As this parts list continues to accumulate widely-expressed “non-muscle” proteins, the hypothesis emerges that the early steps of the sarcomere assembly program in muscle cells are largely re-purposed from the contractile assembly programs of ancient, primitive cells. This hypothesis is also in line with well-known evidence that sarcomeres arise directly from actin arc-like structures resembling those in non-muscle cells^6,7^. In light of this, we propose that similar roles, during CM sarcomere assembly, might exist for other “non-muscle” paralogs of muscle proteins.

## METHODS

### Cell culture

<u>**H**</u>uman <u>**i**</u>nduced pluripotent stem cell–derived <u>**c**</u>ardiac <u>**m**</u>yocytes (hiCMs) were obtained from Fujifilm Cellular Dynamics (iCell Cardiomyocytes2; cat# CMM-100-012-000.5) and cultured in accordance with the manufacturer’s instructions. Briefly, the cells were thawed and seeded at a density of 50,000 cells per well in gelatin-coated, 96-well polystyrene plates (EMD Millipore; cat# ES-006-B). After five hours, the plating medium was replaced with Cardiomyocyte Maintenance Medium (Fujifilm Cellular Dynamics; cat# M1003), which was then refreshed every 48 hours. Cultures were maintained at 37°C in a 5% CO_2_ atmosphere.

### hiCMs re-seeding/sarcomere assembly assay

To investigate the effects of various perturbations on cardiac myocyte sarcomere assembly, hiCMs cultured in 96-well plates were rinsed twice with 100 µL warm PBS lacking Ca^2+^ and Mg^2+^ (Gibco cat# 70011044). Sarcomere disassembly was initiated by trypsinization for 2.5 minutes in 40 µL of 0.1% Trypsin-EDTA lacking phenol red (Gibco cat# 15400054), followed by gentle pipetting to detach cells from the substrate. Trypsin was neutralized, and the cells placed into suspension, by the addition of 120 µL of pre-warmed Cardiomyocyte Maintenance Medium. Cell suspensions were then pelleted by centrifugation at 0.2 rcf for 3 minutes. After aspirating off the supernatant, cell pellets were gently resuspended in 100 µL of fresh, pre-warmed Cardiomyocyte Maintenance Medium. Finally, the resuspended cells were carefully dispensed onto 10 µg/mL fibronectin (Corning cat# 354008) pre-coated cover glasses of various types of 35 mm Cellvis plates, including either four-chamber plates (cat# D35C4-20-1.5-N) or 10mm single cover glass plates (cat# D35-10-1.5-N), depending on experimental needs/design.

### hiCMs siRNA-mediated depletion (knockdown) protocol

To test the effects of siRNA-dependent protein depletion, hiCMs were subjected to the following protocol prior to re-seeding: siRNA that was either non-targeting (“siScramble”; cat#D-001910-01-05) or targeted against either human *ACTN1* (“si-*ACTN1* single”; Accel# A-011195-13-0005 or “si-*ACTN1* pool”; Accel# E-011195-00-0005) or *ACTN2* (“si-*ACTN2*”; Accel# A-011196-16-0005) was pre-mixed with Lipofectamine RNAiMAX (Invitrogen; cat# 13778-075) for five minutes in room temperature Optimem (Gibco cat#11058-021), as according to RNAiMAX manufacturer’s instructions. After five minutes, siRNA pre-mixtures were re-suspended in pre-warmed Cardiomyocyte Maintenance Medium and added directly to hiCMs. Protein levels after depletion were assessed via immunoblotting against siRNA-targeted protein, averaged across three separate biological replicate experiments. For each experiment, tubulin levels (TUBA1A, immunoblotted using antibodies against DM1A Thermofisher cat# 62204) were used as a loading control.

### hiCMs transfection protocol

For transfection of hiCMs, cells were subjected to the following protocol prior to re-seeding: Promega ViaFect (cat# E498A) was combined with 100ng plasmid DNA in room-temperature Opti-Mem (Gibco; cat# 11058-021) at a ratio of 0.6uL ViaFect:100ng DNA and allowed to incubate at room-temperature for 20-45 minutes. The ViaFect/DNA mixture was then mixed into cardiomyocyte maintenance medium, added directly to hiCMs, and allowed to incubate for 18-24 hours. Transfected hiCMs were then re-plated according to the standard reseeding assay protocol. Plasmids used for transfection in this study were paxillin-tdEOS (addgene 57653), vinculin-eos3.2 (addgene 66952), ACTN1-mEmerald (available upon request) and ACTN2-mEmerald (available upon request).

### Immunofluorescence protocol

hiCMs were chemically fixed 24 hours after re-reseeding. Fixation was performed by a 20-minute exposure to a room-temperature 4% paraformaldehyde (PFA) in PBS solution. Fixed hiCMs were then permeabilized in a 4% PFA + 1% Triton X-100 solution. Permeabilized hiCMs were then rinsed three times in 1X PBS, then blocked using a 10% bovine serum albumin (BSA) solution suspended in PBS (blocking buffer). After a minimum blocking period of 20 minutes, hiCMs were incubated in blocking buffer + primary antibodies overnight at 4 C. The next morning, hiCMs were rinsed three times with blocking buffer, then incubated in blocking buffer + secondary antibodies at room temperature for one hour. After secondary incubation, hiCMs were washed three times with PBS, then incubated in PBS + DAPI for 20 minutes. After DAPI incubation, hiCMs were washed three times with PBS. Fields of view were selected for imaging using DAPI to avoid imaging bias. Antibodies used in this study were anti-ACTN1 (Invitrogen, PA5-44889), anti-ACTN2 (Sigma, EA-53), anti-paxillin (BD Biosciences, 610051).

### Z-lines/cell and adhesions/cell analysis

For each field of view, a 3-dimensional image stack was acquired and subsequently converted to a 2-dimensional maximum z-projection. For Z-lines/cell analysis, maximum z-projections of ACTN2 stains were automatically binarized, then thresholded via the deep-learning tool, yoU-Net^31^. Z-lines per cell were quantified from yoU-Net-generated binaries via the automatic analysis tool, sarcApp^31^.

For average adhesion size/adhesions per cell analysis, maximum z-projections of paxillin stains (or images of paxillin-eos) were binarized and thresholded manually, then analyzed using FIJI Analyze Particles with a size exclusion criterion of 0.2 um^2. Particles above 7.0 um^2 represented thresholding artifacts and were excluded from adhesion analyses.

### Photo-conversion of paxillin-eos and vinculin-eos

To investigate the effects of ACTN1 depletion on the stability of adhesion proteins, hiCMs subjected to the depletion protocol were transfected with plasmid DNA encoding either photo-convertible paxillin-eos or vinculin-eos one day prior to re-seeding. The next morning, transfected hiCMs were seeded onto four-chamber Cellvis dishes pre-coated with 10 µg/mL human fibronectin and allowed to spread for 18 hours. After the 18-hr spreading period, fields containing transfected cells (i.e., the green/pre-photoactivation channel) were selected for each treatment group and a circular region-of-interest (ROI) was drawn around a fluorescent adhesion. Prior to photo-conversion, a single image was acquired in the red channel for quantification of red channel pre-photo-conversion gray-levels. Then, a brief pulse with a 405-nm laser was employed to photo-convert the pre-drawn ROI, converting a population of fluorescent signal in the adhesion to emit red instead of green. Red fluorescence was then imaged for two hours. To plot fluorescence intensity over time after photo-conversion, red fluorescence signal for each cell was normalized to maximum signal using GraphPad Prism. Decay half-life and mobile fraction were each calculated from non-linear curve-fits assuming one-phase exponential decay from the normalized red fluorescence loss curves of each individual cell.

### Western blot

hiCMs were trypsinized similar to the re-seeding protocol, then pelleted by centrifugation at 0.2 rcf for 3 minutes. After discarding the supernatant, the cell pellet was re-suspended in 400 μL of PBS. Re-suspended pellets were then centrifuged again at 0.3 rcf for 4 minutes. After discarding the supernatant, the remaining pellet was lysed on ice for 45–60 minutes in CellLytic M buffer (Sigma C2978) supplemented with a protease inhibitor cocktail. After lysis, samples were centrifuged at 13,000 rpm for 20 minutes at 4°C. After centrifugation, resulting supernatants were mixed with LDS sample buffer (Life Technologies, NP0007) and sample reducing buffer (Life Technologies, NP0009) before loading onto a 4–12% bis-tris pre-cast gel.

Gel electrophoresis was run at 100 V and proteins were transferred onto 0.45 μm nitrocellulose membranes (Protran NBA085C001EA) via wet transfer at 100 V for 1 hour and 15 minutes in NuPage transfer buffer (Life Technologies, NP0006-1) containing 10% methanol and 0.1% SDS. Following transfer, membranes were blocked for 1 hour at room temperature in a 5% nonfat dry milk solution (Research Products International, M17200-500.0) prepared in Tris-buffered saline with Tween-20 (TBST; 10X TBS from Corning 46-012-CM, Tween-20 from Sigma P7949). Membranes were then incubated overnight at 4°C with primary antibodies diluted in the same 5% milk/TBST mixture. The next morning, membranes were washed in TBST three times, 5 minutes each, then incubated at room temperature for 1 hour with HRP-conjugated secondary antibodies. After three additional 5-minute washes in TBST, membranes were visualized using chemiluminescence imaging. Anti-bodies used in western blots were anti-ACTN1 (Invitrogen, PA5-44889) and anti-TUBA1A (Thermofisher DM1A).

### Microscopy

For Z-lines/cell and adhesions/cell analysis, images were acquired using an instant Structured Illumination Microscopy (iSIM) microscope equipped with a Visitech iSIM system and either a Nikon SR HP Apo TIRF 100x oil immersion objective (model MRD01997, NA=1.49) at 1X zoom or the 60X oil immersion objective. Images were acquired on the iSIM through a Hamamatsu ORCA-Fusion Digital CMOS camera (model C14440-20UP) set to an axial step-size of 0.1 µm. Images were deconvolved using FIJI-integrated, Microvolution software (Cupertino, CA) over 10 iterations set to the Blind method.

For high-resolution images of adhesions, we utilized a Nikon 100X immersion F-oil (refractive index 1.515) objective (NA=1.45) of a Nikon CSU-W1 SoRa microscope in “SoRa mode” (i.e., at 240X). Post-acquisition, images were deconvolved over 10 iterations using the Blind method.

### Sources of Funding

This work was supported by a grant from the National Institute of General Medical Sciences (R35 GM125028) to DTB and a graduate student fellowship from the American Heart Association (836090) to JBH.

